# The Bio::Phylo libraries for phylogenetic data analysis, version 2.0

**DOI:** 10.1101/211334

**Authors:** Rutger A. Vos, Hannes Hettling

**Affiliations:** Naturalis Biodiversity Center, Leiden, P.O. Box 9517, 2300RA, The Netherlands; Institute of Biology Leiden, Leiden University, Leiden, P.O. Box 9500, 2300RA, The Netherlands

## Abstract

**Motivation:** Phylogenetic analysis is a broad and expanding field that requires versatile programming toolkits to manage the various data types, file formats, and needs for scalability, simulation, visualization, and data exploration.

**Results:** We present version 2.0 of the Bio::Phylo libraries for phylogenetic data analysis. This new release represents a rewrite of the architecture, allowing for extensions that improve speed and persistence, as well as increased functionality in terms of analysis, data reading and writing, and visualization.

**Availability:** The package is released as open source software under the same terms as Perl itself and available from the comprehensive Perl archive network as well as directly from the source code repository.

**Supplementary information:** Supplementary data are available as `doi:10.5281/zenodo.1039210`

## 1 Introduction

Phylogenetic data is multi-faceted, encompassing morphological observations, molecular sequences, biological taxa, and tree and network topologies, and is ubiquitous in a variety of research fields including comparative genomics, systematics, evolution, biodiversity research, and ecology. The types of operations that are performed on phylogenetic data range from data cleaning, integration, and conversions, to simulation, character analysis and inference, and visualization. Flexible programming toolkits that operate on phylogenetic data and aid in scripting these operations are therefore very useful and available in a variety of programming languages. Bio::Phylo (Vos *et al.*, 2011) is the most versatile toolkit for handling phylogenetic data in the Perl programming language.

Conceived about ten years ago (Vos, 2006, 2017), Bio::Phylo has been under ongoing development ever since. Numerous people have contributed to it, by writing code – as volunteers and Google Summer of Code students – by submitting bug reports, and by providing inspiration and impetus in larger collaborative networks (e.g. Stoltzfus *et al.* (2010), Koureas *et al.* (2016b), Koureas *et al.* (2016a)) and hackathons (e.g. Lapp *et al.* (2007), Katayama *et al.* (2010), Katayama *et al.* (2011), Katayama *et al.* (2013), Stoltzfus *et al.* (2013), Katayama *et al.* (2014), Vos *et al.* (2014)). In the process, additional requirements surfaced, which have been addressed by a rewrite of the internal architecture and the addition of functional modules. These have been integrated in a version 2.0.0 release, which we present here.

## 2 Design

In ‘object-oriented’ programming, the different concepts within the problem domain that the software addresses are modelled as different classes (e.g. a class to model phylogenetic trees, and one to model multiple sequence alignments) that inherit some of their functionality from other such classes. For example, a class that represents the blueprint for a multiple sequence alignment might inherit certain generalized attributes of character state matrices (e.g. the number of characters and taxa) from a ‘super-class’. In turn, this multiple sequence alignment class can be inherited from to create a blueprint for a more specialized class, for example one that models amino-acid alignments.

The object-oriented programming paradigm allows other programmers to extend the functionality of toolkits by inheriting from classes to address additional use cases, for example by improving performance in various ways. Bio::Phylo has always allowed for this, but the version 1.0 design made this relatively cumbersome because a lot of functionality would have to be re-implemented for any newly created class to integrate well in the rest of the toolkit. In the new design, each class has been reworked into two separate modules, one that performs the actual changes to any data associated with the class, and one that performs all the other user-friendly operations that do not directly affect the data. With this new design, a new class that only re-implements the changes to the data can re-use all the other operations, and thus be much more compact and easily written. In addition, the integration of such new classes has been made far more flexible (by adopting, in the parlance of software design patterns (Gamma *et al.*, 1995), the ‘Factory’ pattern throughout).

The usefulness of this change of design is demonstrated by two optional extension packages that can be installed alongside Bio::Phylo. One of them, Bio::PhyloXS, serves as a drop-in replacement of the core data objects in Bio::Phylo, re-implemented in the C programming language. Because C is a very fast, compiled language, having some of the core functionality (e.g. setting and fetching data properties) leads to significant speed increases: a simple benchmark test where a small tree topology is constructed and then traversed executes about 700% faster using this extension. (Note, however, that bindings between C and Perl are fraught with challenges due to the complexity of the Perl API and the differences in memory management on different architectures. Hence, this application should currently be considered ’experimental’.) The second optional extension, Bio::Phylo::Forest::DBTree, allows very large trees to be stored into, and accessed from, a simple SQLite database. This means that these trees never have to be loaded in working memory, and that, once stored in the database, there is no "file reading" step. The typical application for this is to store static, immutable trees. To demonstrate this, we make available database files that were created by indexing the releases listed in Table 1.

**Table 1.** Large, published phylogenies made available as database files.

| Project | Database files |
| --- | --- |
| PhyloTree (van Oven and Kayser, 2009) | 10.6084/m9.figshare.4620757.v1 |
| D-Place (Kirby *et al.*, 2016) | 10.6084/m9.figshare.4620217.v1 |
| NCBI taxonomy (Federhen, 2011) | 10.6084/m9.figshare.4620733.v1 |
| Greengenes (DeSantis *et al.*, 2006) | 10.6084/m9.figshare.4620214.v1 |

## 3 Data input and output

In addition to the file formats already supported in previous releases of Bio::Phylo, several new formats are now read and/or written. Specifically, PhyloXML (Han and Zmasek, 2009), New Hampshire eXtended (NHX, Zmasek and Eddy (2001)), the extensions added to the NEXUS standard by the programs TreeAnnotator and Figtree (Rambaut, 2007), and a ‘badgerfish’ mapping of NeXML (Vos *et al.*, 2012) to JSON can now both be read and written. In addition, several response formats from web services (namely, the DarwinCore archive format returned by GBIF (Baker *et al.*, 2014), the response format from the TaxoSaurus.org service (Stoltzfus *et al.*, 2013), and the response documents of uBio.org) as well as FASTQ (Cock *et al.*, 2009) files can be read. Data files for Hennig86 (Farris, 1988) and for the haplotype ‘Network’ program (fluxus-engineering.com) can be written.

Support for RDF is experimental: RDF/XML triples can be written by way of a two-step process that first exports data as NeXML and then transforms this to statements that obtain terms from CDAO (Prosdocimi *et al.*, 2009); this is the same approach and uses the same transformation stylesheet as TreeBASE (Piel *et al.*, 2009)). RDF graphs can also be ‘read’ in the sense that a graph can be loaded and interrogated with SPARQL queries to extract statements that map onto Bio::Phylo’s object model. At present, this means that annotated taxa and tree topologies can be extracted from CDAO/RDF, but character data cannot.

## 4 Analysis, modification, and simulation

From its first versions onwards, Bio::Phylo has implemented numerous algorithms for computing topological indices, simulating new topologies, and adjusting them algorithmically. Some of the simulation methods were developed and discussed by Hartmann *et al.* (2010), while some of the topology index algorithms were explored by Martyn *et al.* (2012). To the existing topology indices, some additional methods were added: the Euclidean distance between trees (see Kuhner and Felsenstein (1994)) and the number of ‘cherries’ in a given topology (see McKenzie and Steel (2000)) cannow becomputed. Tothealgorithms thatadjusttopologies have been added methods to obtain ultrametric trees using the mean path lengths method of Britton *et al.* (2002), by making node ages proportional to clade size (as per Grafen (1989)), and using an implementation of Stadler’s algorithm for computing the relative order of speciation or coalescence events on a given phylogeny (Gernhard *et al.*, 2006).

By implementing a bridge between Bio::Phylo’s library code and the R programming language, additional functionality has become accessible. So far, this has been wrapped in methods to estimate the parameters of the birth/death process and use these to simulate replicates of the input tree (using ‘ape’, Paradis *et al.* (2004)); and methods to estimate the parameters of state transition models for binary characters and DNA sequences (using ‘phangorn’, Schliep (2010); ‘phylosim’, Sipos *et al.* (2011); and ‘phytools’, Revell (2012)). To encapsulate these state transition models, a class hierarchy has been implemented that represents some of the common substitution models (i.e. JC69, HKY85, GTR, F81, K80) such that they can be serialized to the syntax of commonly used programs for phylogenetic inference. In addition, this facility to select substitution models has been combined with a (tree-based) sequence simulator to allow multiple sequence alignments to be replicated.

## 5 Visualization

The ability to visualize phylogenies as rooted, rectangular cladograms and phylograms in vector formats (SVG being best supported) has always existed in Bio::Phylo. This included functionality to paint branches, add pie charts to nodes, collapse clades as triangles, and influence styling (such as branch thickness; fonts and their size, weight, and style). To this has been added in v2.0.0 the ability to draw unrooted and radial tree projections, andtheoptiontomarkup higher taxa in labeledbraces (straight or arched, depending on projection). Figure 1 demonstrates some of this new functionality.

**Fig. 1.**
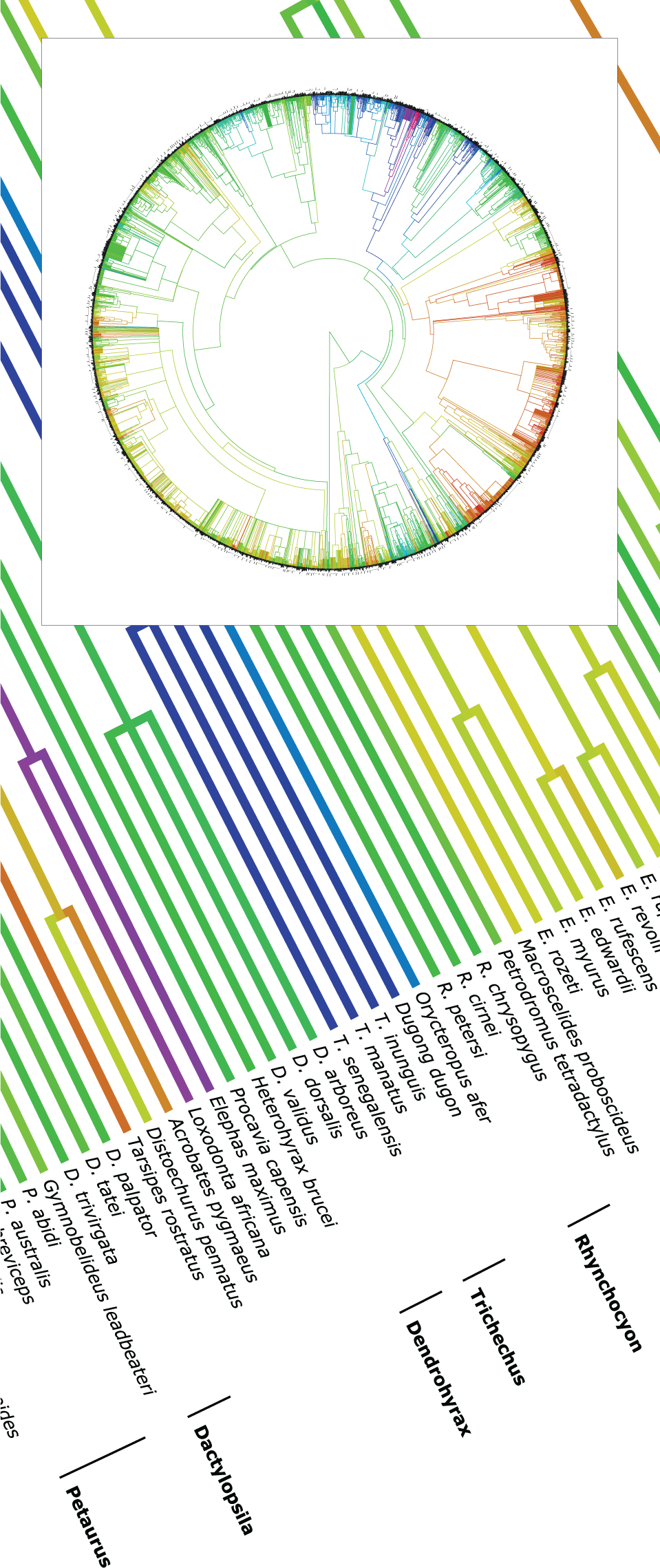
Mammal supertree (Bininda-Emonds et al., 2007) decorated with log-transformed body mass (Jones et al., 2009) as branch colors and monophyletic genera as labeled braces; detail and full tree (inset). Image produced as per the workflow described on http://rvosa.github.io/bio-phylo/doc/examples/integration/

## 6 Impact and re-use

As of time of writing (October 2017), the first version of Bio::Phylo has been cited 35 times. Among these, the papers with the highest impact used Bio::Phylo in analyses in phylogenomics and comparative genomics (e.g. see Roure *et al.* (2012); Hayward *et al.* (2013); De Smet *et al.* (2013)).

Several infrastructural projects depend on Bio::Phylo. These include the BioPerl project (Stajich *et al.*, 2002), which uses it for reading and writing NeXML. This usage is promoted by the interface compatibility between the two projects: the core data objects of Bio::Phylo, i.e. trees, tree nodes, sequence alignments, and so on, can be used directly in BioPerl. Also, the TreeBASE project (Piel *et al.*, 2009) uses Bio::Phylo for certain server-side maintenance tasks, and the SUPERSMART project (Antonelli *et al.*, 2017), as well as the BioVeL services that expose it (Hardisty *et al.*, 2016), are deeply integrated with Bio::Phylo (and drove some of the development of new functionality).

In addition, several single-purpose programs and pipelines use Bio::Phylo. These include the Monophylizer web service for assessing monophyly in gene trees (Mutanen *et al.*, 2016; Pentinsaari *et al.*, 2016); the CopyRighter tool for improving the accuracy of microbial community profiles through lineage-specific gene copy number correction (Angly *et al.*, 2014); and the PhyloMatch pipeline to discover highly phylogenetically informative genes in sequenced genomes (Ramazzotti *et al.*, 2012).

## 7 Availability

All revisions of the source code are available from the source code repository at http://github.com/rvosa/bio-phylo. The latest stable release version, which over time may fall behind the latest source code revision, is available from the Comprehensive Perl Archive Network (CPAN) at http://search.cpan.org/dist/Bio-Phylo. Accompanying this publication is a uniquely identifiable release, stamped with a Digital Object Identifier (DOI) issued by Zenodo.org: doi:10.5281/zenodo.1039210.

As is a common convention in Perl software releases, a dual licensing scheme applies to Bio::Phylo – both the Artistic License (https://github.com/rvosa/bio-phylo/blob/master/COPYING) as well as the GNU General Public License (https://github.com/rvosa/bio-phylo/blob/master/LICENSE) applies. This is generally interpreted to mean that you are free to choose whichever of these licenses fits best with your own project, should you want to reuse all (or part) of Bio::Phylo. This is certainly the spirit: feel free to use these libraries however you see fit. No warranties.

## Acknowledgements

The following people have contributed code to the project: Florent Angly, Jason Caravas, Klaas Hartmann, Mark A. Jensen, Moritz Lenz, Chase Miller, Aki Mimoto, and Jan Willem Wijnands. The following people have provided feedback through bug reports and reviews: Denis Baurain, Chris Fields, Shlomi Fish, Jean-Marc Frigerio, Andreas J. König, Hilmar Lapp, Nicolas Lenfant, Sébastien Moretti, Slaven Rezić, Seiler, and scorpio17. Mannis van Oven has been very helpful in providing the data dumps of the PhyloTree.org project that were indexed as database files (in Table 1). The principal investigators of the labs in which RAV has done his research have all allowed and encouraged the development of Bio::Phylo. These are Arne Mooers, Wayne Maddison, and Mark Pagel, and now Naturalis Biodiversity Center. We are grateful to all these people for their support.

